# Improved taxonomic annotation of Archaea communities using LotuS2, the Genome Taxonomy Database and RNAseq data

**DOI:** 10.1101/2023.08.21.554127

**Authors:** Alastair Grant, Abdullah Aleidan, Charli S. Davies, Solomon C. Udochi, Joachim Fritscher, Mohammad Bahram, Falk Hildebrand

## Abstract

Metabarcoding is increasingly used to uncover diversity and characterise communities of Archaea In various habitats, but taxonomic annotation of their sequences remains more challenging than for bacteria. Fewer reference sequences are available; widely used databases do not reflect recent revisions of higher level archaeal taxonomy and a substantial fraction of their phylogenetic diversity remains to be fully characterised. We address these gaps with a systematic and tractable approach based around the Genome Taxonomy Database (GTDB). GTDB provides a standardized taxonomy with normalized ranks based on protein coding genes, allowing us to identify and remove incongruent SSU sequences. We then use this in combination with the eukaryote PR2 database to annotate a collection of near full length rRNA sequences and the Archaea SSU sequences in SILVA, creating a new reference database, KSGP (***K***arst, ***S***ilva, ***G***TDB and **P**R2). GTDB SSUs alone provides a small improvement in annotation of an example marine Archaea OTU data set over standardized SSU databases such as SILVA and Greengenes2, while KSGP increases Class and Order assignments by 145% and 280% respectively and is likely to provide some improvement in annotation of bacterial sequences too.

We make the KSGP database and a cleaned and deduplicated subset of GTDB SSU sequences available at ksgp.earlham.ac.uk; integrate them into a metabarcoding pipeline, LotuS2 and outline rapid and robust strategies to generate a set of annotated Archaea OTUs and to determine the proportion of Archaea sequences in metatranscriptomic data. We also demonstrate simple tools to visualise the completeness of database coverage and outline strategies to further understand poorly characterised components of the archaeal community which will be equally applicable to bacteria.

## Introduction

Archaea were initially viewed as extremophiles inhabiting hot springs, salt lakes and acid mine drainage, but are now recognised as ubiquitous and diverse in benign environments such as soil, wetland, oligotrophic oceans and the animal gut, where they can make up a significant fraction of biomass and carry out important biogeochemical processes (Adam, Borrel, Brochier-Armanet, & Gribaldo, 2017; Bahram, Anslan, Hildebrand, Bork, & Tedersoo, 2019; Bahram et al., 2022; Gribaldo & Brochier-Armanet, 2006; Karner, DeLong, & Karl, 2001; Martin-Pozas et al., 2020; Rinke et al., 2021; Tahon, Geesink, & Ettema, 2021; Woese & Fox, 1977; Youngblut et al., 2021). However metabarcoding methods for Archaea, particularly taxonomic annotation, are not as well established as for bacterial communities (Abellan-Schneyder et al., 2021; Bahram et al., 2019; Belilla et al., 2019; Djurhuus et al., 2017; George et al., 2019; Hugoni et al., 2018; Wurzbacher et al., 2017). Important reference databases such as SILVA (Pruesse et al., 2007), Greengenes (DeSantis et al., 2006; D. McDonald et al., 2012a) and RDP (Cole et al., 2009) contain archaeal sequences in much lower numbers than bacteria. These are often incompletely annotated, owing to lack of reference genomes, difficulty of culturing many Archaea and archaeal taxonomy lagging behind the much more well-defined bacterial taxonomy (Adam et al., 2017; Parks et al., 2020). NCBI RefSeq provides reference SSU sequences, but these also suffer from taxonomic mis-annotations (Smith, Glendinning, Walker, & Watson, 2022). Edgar (2018a) estimated that ∼17% of taxonomic annotations in the SILVA and Greengenes databases are incorrect because of shortfalls with automatic taxonomic assignment and it is unlikely that annotations for the much less well studied Archaea will be more reliable than this average. In addition, substantial discordance between archaeal phylogeny and the current NCBI taxonomy was reported recently (Parks et al., 2020) and “universal” 16S rRNA gene primers are often biased against Archaea, especially newly discovered, lineages (Bahram et al., 2019; Eloe-Fadrosh, Ivanova, Woyke, & Kyrpides, 2016; Karst et al., 2018).

A route to improve this situation is provided by the standardised archaeal taxonomy based on a phylogeny constructed from conserved single copy marker genes in the prokaryote Genome Taxonomy Data Base project (hereafter GTDB; Parks et al., 2020; Rinke et al., 2021). This assigns all available Archaea genomes, including metagenomic assembled genomes (MAGs), to “species clusters” and normalises taxonomic ranks of these at levels from genus to phylum based on coding gene sequence divergence. 16S rRNA gene sequences (henceforth “16S sequences”) have been identified in each genome, but were not used in phylogeny reconstruction. This standardizing taxonomic approach proposed 16 phyla of Archaea, uniting Thaumarchaeota, Aigarchaeota, Crenarchaeota and Korarchaeota into the phylum Thermoproteota and splitting the former Euryarchaeota into five separate phyla (Hadarchaeota; Hydrothermarchaeota; Methanobacteriota; Thermoplasmatota and Halobacteriota).

Here we demonstrate how GTDB can be used both on its own and in conjunction with collections of near full length SSU sequences and PCR amplicons to obtain improved and consistent taxonomic annotations of Archaea sequences in both metabarcoding and RNASeq data of Archaea.

## Materials and methods

### Databases used

The results reported here are based on GTDB version 214.0; the Sativa subset of GTDB release 207 (Lundin & Andersson, 2021, hereafter *Sativa*); SILVA version 138.1 (Quast et al., 2012); Greengenes 13.5 (Daniel McDonald et al., 2012b) Greengenes2 2022.10 (Daniel McDonald et al., 2023) and PR2 5.0.0 (Guillou et al., 2012; Vaulot et al., 2022).

### Cleaning the GTDB 16S sequence database

Taxonomic assignments in GTDB are underpinned by a phylogeny based on protein coding genes derived from genome sequences of isolated strains and MAGs (Parks et al., 2020; Parks et al., 2018). 16S sequences are available for these genomes, but were not used in phylogeny construction. Our initial analyses showed that some GTDB 16S sequences annotated as Archaea were, in fact, Bacteria and vice versa so some database cleaning is required. Others have also noted misclassifications and inverted orientations of GTDB 16S sequences (Kozlov, Zhang, Yilmaz, Glöckner, & Stamatakis, 2016), although from release 08-RS214 of GTDB, all sequences are in 5’ to 3’ orientation (Parks, 2023). A subset of GTDB has previously been produced by stringent filtering of short and low quality sequences followed by semi-automated removal of mis-classified sequences (Lundin & Andersson, 2021, hereafter SATIVA). The GTDB taxonomy is independent of the 16S sequences, so we were able to use the *Classify* command of RDPtools and the classify and training set No. 18 (RDP, 2020; Wang, Garrity, Tiedje, & Cole, 2007), to assess whether 16S sequence differences were congruent with the taxonomy without having to base this on a guide tree constructed from the sequences themselves. We removed 5000 sequences misclassified at domain level by this process (see results for details) then removed duplicate sequences with the *rm_dupseq* command of RDPtools Classify. The RDP *Loot* (leave one out) command was then used to identify a further 47 sequences which still appeared to be incorrectly assigned at Domain level. A list of the sequences removed by these processes is in table S1.

PCR primers targeting archaeal SSU can amplify eukaryote sequences (AG, personal observation; and see results, particularly Fig. 3a), so it is necessary to supplement any prokaryote database with sequences spanning the range of eukaryote diversity likely to be present. Here we use the manually curated PR2 database (Vaulot et al., 2022, available in LotuS2) after removing the prokaryote sequences that it contains. PR2 is orientated towards protists, particularly from the marine environment so alternatives such as SILVA eukaryote sequences may be preferable for data from terrestrial systems. We are using the PR2 database primarily to remove rather than identify eukaryote sequences, so unless otherwise indicated, in the remainder of this paper we use GTDB+ as a shorthand for the cleaned and deduplicated GTDB sequences concatenated with PR2 eukaryote sequences.

### Annotating the Karst et al (2018) database and SILVA Archaea sequences

Karst et al. (2018, ENA accession GCA_900214305) provide an unannotated collection of > 1 million bacterial, archaeal and eukaryote SSU rRNA sequences longer than 1200 bp (hereafter KARST). Sequences annotated as archaeal were extracted from the Ref_NR99 SILVA SSU database (Quast et al., 2012). Taxonomic assignments of both of these databases were made using GTDB+ and SINTAX (Edgar, 2016a) with a probability cut-off of 80%. SINTAX is one of the highest accuracy classifiers (Hung et al., 2022) and the k-mer matching algorithm it uses is much more tolerant of database sequences of varying lengths than are approaches based on phylogenetic tree construction. The SINTAX output file was reformatted into a LotuS2 taxonomy file and used to annotate OTUs in conjunction with a FASTA file containing the corresponding sequences. In the remainder of this paper we refer to the combined database of GTDB, PR2; the annotated Karst sequences and reannotated SILVA Archaea sequences as KSGP (***K***arst, ***S***ilva, ***G***TDB and **P**R2). Subsequent references to SILVA are to version 138.1 with its original annotations, except where explicitly indicated.

### Example metabarcoding and metatranscriptomic data sets

Archaea metabarcoding and RNASeq data were generated for DNA and RNA extracted from estuarine sediments along a gradient ranging from uncontaminated to sites severely impacted by metal pollution, particularly copper. Here we characterise the Archaea OTUs present in the pooled sequencing data to illustrate our methodology. Relationships between microbial communities and the contamination gradient, and the detailed taxonomic composition of these communities, will be examined elsewhere.

### Sample collection, DNA & RNA extraction

A total of 47 samples were collected. RNA and DNA were extracted from three replicates at 11 intertidal locations in estuaries in SW England and one in Breydon water, Norfolk (details in Udochi, 2020). DNA was extracted from a further 11 samples from Breydon water. DNA and RNA were extracted from 1.4 – 2.7g of each sample using the RNeasy PowerSoil Total RNA kit and DNA elution accessory kit (Qiagen, Hilden, Germany) according to an optimised version of the manufacturer’s instructions with an added heat block step (45°C for 15 minutes) prior to the solution being added to the column. Nucleic acids were quantified using Invitrogen Qubit RNA and dsDNA broad range kits (ThermoFisher, Loughborough, UK) measuring fluorescence with either a qPCR machine or a Qubit 4 fluorimeter.

### PCR and sequencing

PCR amplification was carried out using the SSU1ArF/SSU520R primer pair (Bahram et al., 2019) with Illumina sequencing adapters, barcodes and length heterogeneity spacers appended to their 5’ ends (following Caporaso et al., 2012). Cycling parameters were an initial denaturation at 98°C for 10 minutes, followed by 35 cycles of denaturation at 98°C for 30s; annealing at 50°C 30s; extension at 72°C for 30s and a final extension at 72°C for 5 minutes. Sequencing was carried out using a single Illumina Novaseq SP flow cell with 250 bp paired end reads, in combination with PCR generated libraries for amplicons of similar length for several other target genes from the same sites. A PCR free genomic library of a saltmarsh grass was added to the flow cell, rather than the PhiX control library normally used to ensure balanced fluorescence across channels when sequencing low diversity libraries. These sequences were removed based on presence of a unique i7 index, whereas the PCR products did not contain i7 indices.

### Total RNA library prep and sequencing

Sequencing libraries were prepared from total RNA using a Corall total RNA-seq library preparation kit (Lexogen, Vienna, Austria) following the manufacturer’s instructions. Sequencing was carried out using a single Illumina MiSeq nano flow cell, with 250 bp paired end reads.

### Bioinformatics

The bulk of analyses were carried out using the LotuS2 pipeline (Ozkurt et al., 2022). This allowed us to rapidly carry out representative examples of the main options for this without the need to run multiple different pipelines. Analysis of the data, including OTU and tree construction and taxonomic annotation with an individual database, required a single command and was completed in under 2 hours on a 64 core Intel Xeon computer. Sequences were demultiplexed and cleaned (Bedarf et al., 2021; Puente-Sanchez, Aguirre, & Parro, 2016); OTUs were clustered at 97% similarity using UPARSE (Edgar, 2013), CD-Hit (Fu, Niu, Zhu, Wu, & Li, 2012) and Swarm (Mahé et al., 2021); zero radius OTUs (zOTUs) were identified using UNoise (Edgar, 2016b) and reads mapped to OTUs/zOTUs using Minimap2 (Li, 2018). Database coverage was assessed by finding the best single match to each OTU using the UBLAST option of USEARCH (Edgar, 2010, with an E value of 0.05) in SILVA, Greengenes, Greengenes2, GTDB, SATIVA, KARST and KSGP. Matches in the NCBI nt database as at 14^th^ July 2023 were found using the BlastN executable version 2.14.0 (Camacho et al., 2009) with an E-value cut-off of 0.05 and a query coverage cut-off of 50%. BlastN on individual sequences of particular interest was carried out via the NCBI Blast web interface. Database coverage was visualised by plotting sequence similarities of the best match in descending order; coverage of pairs of databases was compared using scatter plots of these similarities with density contours superimposed.

Taxonomic assignments from species to phylum were carried out using USEARCH_global searches against the SILVA, GTDB+ and KSGP databases followed by the LCA v0.24 algorithm within LotuS2, using default parameters. Best matches for individual RNAseq sequences were found using USEARCH_global within Vsearch using GTDB+ and KSGP with an identity cut-off of ≥0.65.

Data handling, graph plotting and statistical analyses were carried out using R version 4.2.1 and the Phyloseq, ggplot2 and ggtree libraries (McMurdie & Holmes, 2013; R Core Team, 2022; Wickham, 2016; Yu, 2020). Subsetting and interconversion of fasta and fastq files was carried out using SeqKit (Shen, Le, Li, & Hu, 2016). Sequencing reads are deposited at ENA (project accession PRJEB65254).

## Results

### Congruence of 16S sequences with GTDB taxonomy based on coding genes

GTDB 214 contains 708,813 16S sequences (including duplicates from the same genome). Of these, 6,370 are annotated as Archaea by GTDB. The RDP classifier assigned 124 of these as bacterial sequences, at a probability > 0.8 for all but 5, and one as eukaryote with low confidence, a misclassification rate of 2%. Many of these are MAGs, and in a number of cases the incongruent sequences are assigned by RDP to bacteria that are characteristic of the same environment, suggesting that they often represent contamination of MAGs with 16S sequences from co-occurring organisms. In addition, 4,875 of the 16S sequences (0.7%) annotated as Bacteria in GTDB were identified as misclassified at domain level. In contrast to Archaea, many of these are genome sequences of cultivated strains of human commensals/pathogens such as *E. coli*, *Klebsiella* and *Acinetobacter*. RDP classifies the great majority of these incongruent SSU sequences as Archaea, but RDP contains only a very small number of eukaryote sequences. BlastN searches against NCBI nt and PR2 identify that the best matching sequences to these are from eukaryotes in over 80% of cases, with plant plastids; fungi and mammals being particularly common, suggesting contamination with non-target material during strain isolation, DNA extraction or sequencing library preparation. After removing sequences misclassified at domain level and deduplicating the remainder, 186,215 unique sequences remain (details of sequences removed are given in Supplementary Table S1).

### Amplicon sequencing success and methods of OTU construction

PCR was unsuccessful for one replicate at each of two sites. 20 million sequences from the remaining 45 estuarine samples were used for OTU construction and a further 4 million low quality reads were assigned to these OTUs (see Ozkurt et al., 2022 for more details of low quality read recovery in LotuS2). Excluding a small number of samples where amplification was poor (n=5, < 10,000 reads), there were between 67,000 and 1.3 million reads per sample (median 600,000). UNoise clustered 25,188 zero radius OTUs (zOTUs, in principle equivalent to amplicon sequencing variants, ASVs), compared with 15,179 OTUs generated by UPARSE. CD-Hit and Swarm generated OTU numbers between these extremes (data not shown). In total, 52% of zOTUs were 100% identical to individual UPARSE OTUs, while 94% had at least 97% sequence similarity. The most abundant three OTUs represented 11, 6 and 5% of reads assigned to OTUs, while the most abundant zOTUs only represented between 0.83 and 0.7% of assigned reads, indicating that as expected UPARSE is “lumping” individual zOTUs together. Here our focus is on higher level classifications, so in the remainder of this paper, we use OTUs generated using UPARSE, but recognise that for some purposes it will be helpful to use ASVs/ zOTUs to examine the relative abundance and distribution of the strains (or closely related species, c.f. Edgar, 2018b) that are grouped together into these OTUs (see Box 1).

### Database coverage

We can consider the OTUs to be scattered throughout a hypervolume with three dimensions for each position in the sequence (ignoring indels). The number of OTUs with a match in a database indicates the proportion of this hypervolume that is covered by that database. This varies between 75.3% and 77.3% for the Greengenes, Greengenes2, GTDB+, SILVA, KARST and KSGP databases, with a slightly higher proportion for NCBI nt (table 1; Fig. 1, light green, dark green, orange, brown, blue, black and purple lines respectively). OTUs without a match may represent uncharacterised phylogenetic diversity or PCR/sequencing artefacts. The number of unique reference sequences that are the closest match to an OTU in our data indicates how densely database sequences are distributed within this hypervolume, varying from 618 for Greengenes2 to 4,536 in KARST and 4,945 in KSGP (table 1). Matches in NCBI nt fall between these extremes, with 2,947 unique accessions.

**Figure 1.**
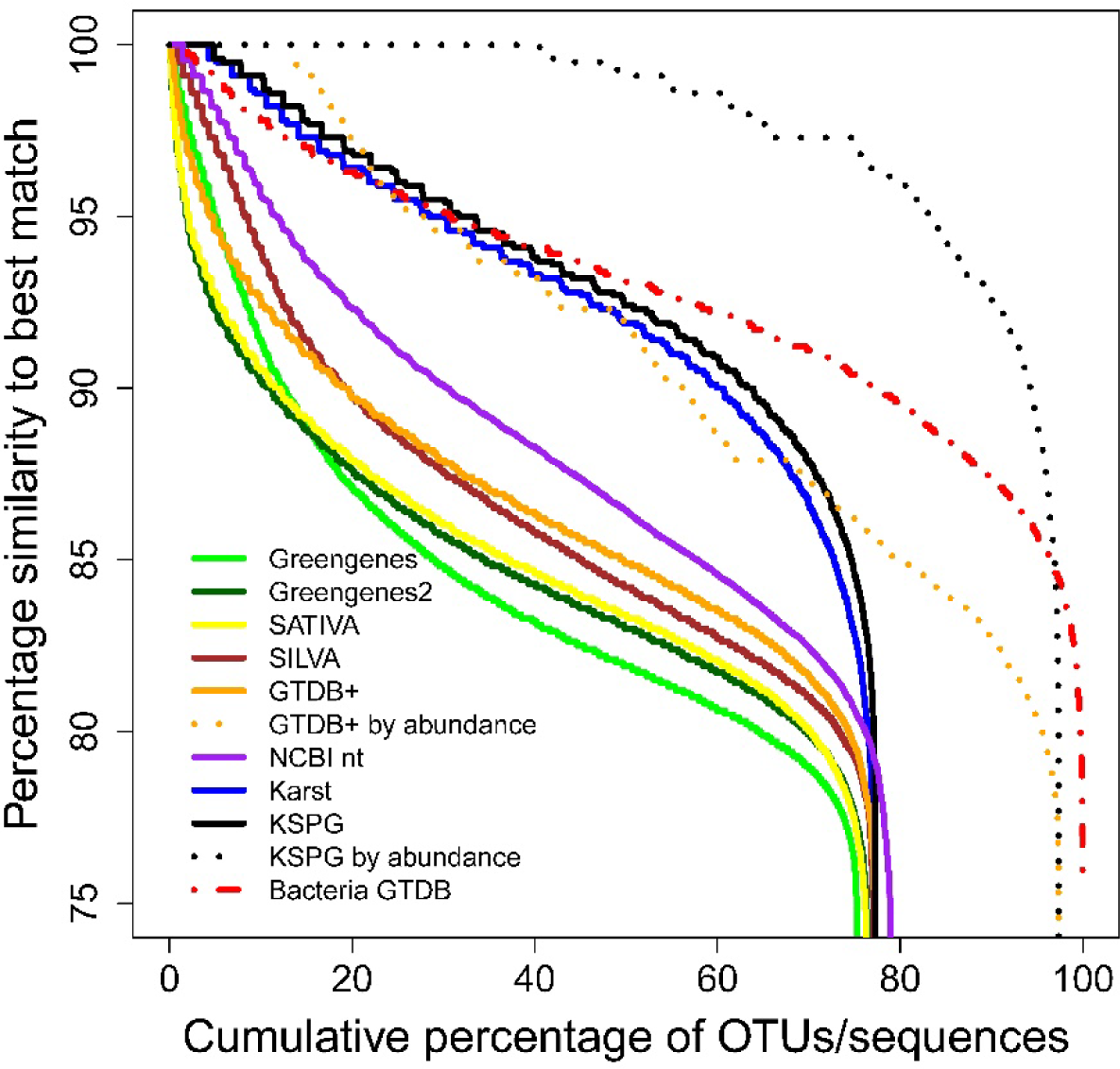
Database coverages. Percentage similarity of all individual OTUs to best hit in Greengenes, Greengenes2, Sativa, GTDB+, SILVA, NCBI nt, Karst and KSGP sorted in descending order and plotted against cumulative OTU rank (solid lines) or cumulative OTU abundance (dotted lines). Data for bacterial OTUs from the same environmental samples are included (red dashed line) to illustrate the current shortcomings in archaeal taxonomic annotations.

**Table 1.** Coverage of Archaea OTUs by the different databases as indicated by the percentage of OTUs for which a Blast match was found and the number of unique accessions in the database represented among these hits.

| Database | Percentage of OTUs with a Blast hit | Number of unique accessions with highest similarity to an OTU |
| --- | --- | --- |
| KSGP | 77.3 | 4945 |
| KARST | 77.1 | 4536 |
| NCBI nt | 83.3 | 2947 |
| SILVA | 77.1 | 1571 |
| Greengenes | 75.3 | 1134 |
| GTDB+ | 77.2 | 1026 |
| SATIVA subset of GTDB | 76.3 | 666 |
| Greengenes2 | 76.5 | 618 |

Cumulative plots of similarities to the closest match (Fig. 1) provide more detail on the distribution of database sequences. For example, SILVA has more matches at high similarity (>90%) than does GTDB+, but GTDB+ performs slightly better at lower similarities indicating that it captures a greater proportion of overall phylogenetic diversity than SILVA. If phylogenetic coverage is constant, we would expect the number of unique reference sequences to be greater for larger databases, but this number is lowest for the largest database, Greengenes2. By contrast, it is closely correlated with the proportion of OTUs with matches at similarities of around 90% indicating the extent to which each database captures phylogenetic diversity at Class and Order level. Remarkably, there are even more high similarity matches in the >1 million SSU sequences provided by Karst et al. (2018) blue line in Fig 1) than in NCBI nt (purple), with the full KSGP database (black) performing only slightly better than this. KARST contains 61,266 archaeal sequences, which cluster into 3,410 OTUs at 97% similarity, but the authors note that many of these have relatively low similarity to previously detected SSU sequences. The bulk of these novel sequences are derived from “sediments”, a category that includes seven estuarine and marine samples and five from freshwater (their Figure 2 and their tables S2 and S5). The inclusion in KSGP of the KARST sequences means that KSGP provides much better coverage than all other databases with the exception of the small number of OTUs which have a match only in NCBI nt. The SATIVA, Greengenes and Greengenes2 databases (yellow and green lines in Fig. 1) perform more poorly than either GTDB+ or SILVA. These are therefore not used in the remainder of our analyses but a more detailed assessment of the performance of the Greengenes2 database on both Archaea and Bacteria is given in the supplementary material, particularly Fig. S1.

When plotted by cumulative sequence abundance rather than OTU rank order (orange and black dotted lines in Fig. 1), over 95% of individual Archaea *sequences* have a match in GTDB+ at ≥80% and in KSGP at >90% similarity, showing that common OTUs are more likely to be successfully annotated than rare ones. By contrast, bacterial OTUs amplified from the same sediments almost all have matches in GTDB+ at higher sequence similarity than do the Archaea OTUs (Fig 1, red dashed line), illustrating the much greater completeness with which bacterial phylogenetic diversity has been characterised.

Figure 2 provides more detailed comparison of taxonomic coverage between the GTDB+ and both SILVA and KSGP. The modal similarity with the best match in GTDB+ is 85%, while similarities with the best match in KSGP have modes at both 95% and ≥99%, showing that the KARST sequences match a substantial fraction of the phylogenetic diversity in our samples at family/genus and species level respectively. SILVA contains more high similarity matches than GTDB+ (Fig. 2b), but the modal similarity in SILVA is approximately 2% *lower*, and the peak of the contour plot is, in consequence, located slightly below the line indicating equal similarity. The fact that many of our OTUs have closer matches in KARST than in the much larger nt database, and the large number of ≥99% matches in particular, will in part reflect the origin of many sequences in KARST from similar environments to our OTUs. But the change in the main mode of the distribution from below 85% for SILVA to 95% for KSPG and the positions of the LOWESS fits to the data suggest that the SSU1ArF/SSU520R primer pair successfully amplifies a wider range of Archaea taxonomic diversity than older PCR primers used to amplify the bulk of environmental Archaea sequences in SILVA.

**Figure 2.**
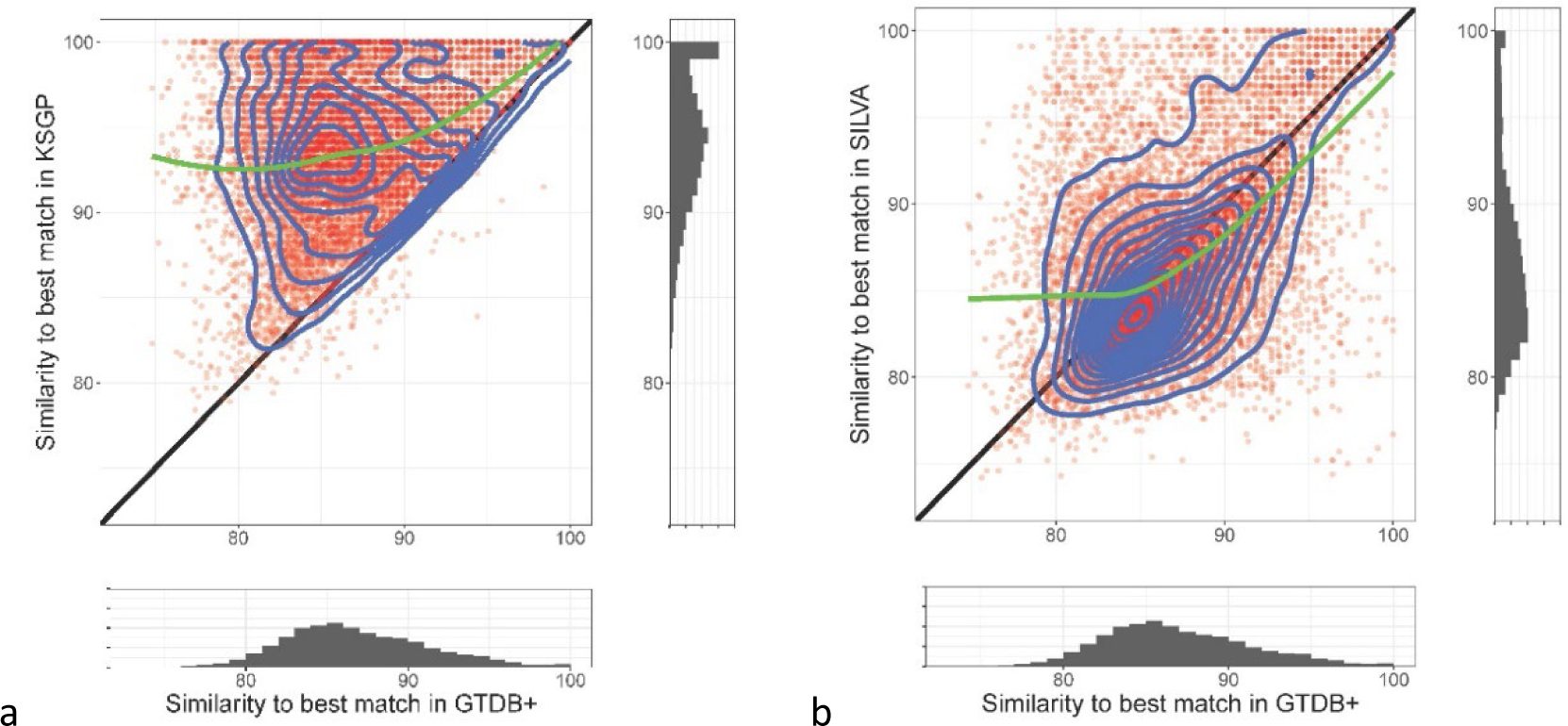
a) Similarity of the best match in KSGP (y-axis) against similarity of the best match in GTDB+ (x-axis). The scatter diagram has density contours superimposed, with marginal histograms depicting density of matches against single databases. Straight line indicates identical similarity in both databases; green line is Loess fit to the data. b) Best matches in SILVA (y-axis) plotted against those in GTDB+. Other details as (a).

### OTUs without database matches

11,442 OTUs (75.4%) are assigned to Archaea and 1.7% eukaryotes using the LotuS2 LCA taxonomy assignments to KSGP. 13 of the 1,000 commonest OTUs were identified as eukaryotes, with OTU 51 being the most abundant. Of the remainder, 22.8% do not have matches in KSGP; 0.1% have good matches to sequences whose domain level taxonomy was not resolved by SINTAX and 0.06% did not have their taxonomy resolved by the LCA analysis. The OTUs without database matches could represent novel archaeal phylogenetic diversity not currently encompassed by the currently used databases; sequences from organisms other than Archaea and/or sequencing/PCR artefacts. The location of unannotated sequences on a maximum likelihood tree of OTUs (Fig. 3a) helps to clarify this question. The great majority of these OTU sequences are rare, none are within the most common 1,000 OTUs. The bulk are located on long branches, forming a large group just below the centre of Fig. 3a and several smaller groups. This suggests that the great majority of our *de novo* generated OTUs that had no match in any of the databases represent sequencing errors or PCR artefacts. Removing these OTUs shifts the cumulative similarity curves rightwards, but has little impact on overall conclusions (data not shown).

**Figure 3.**
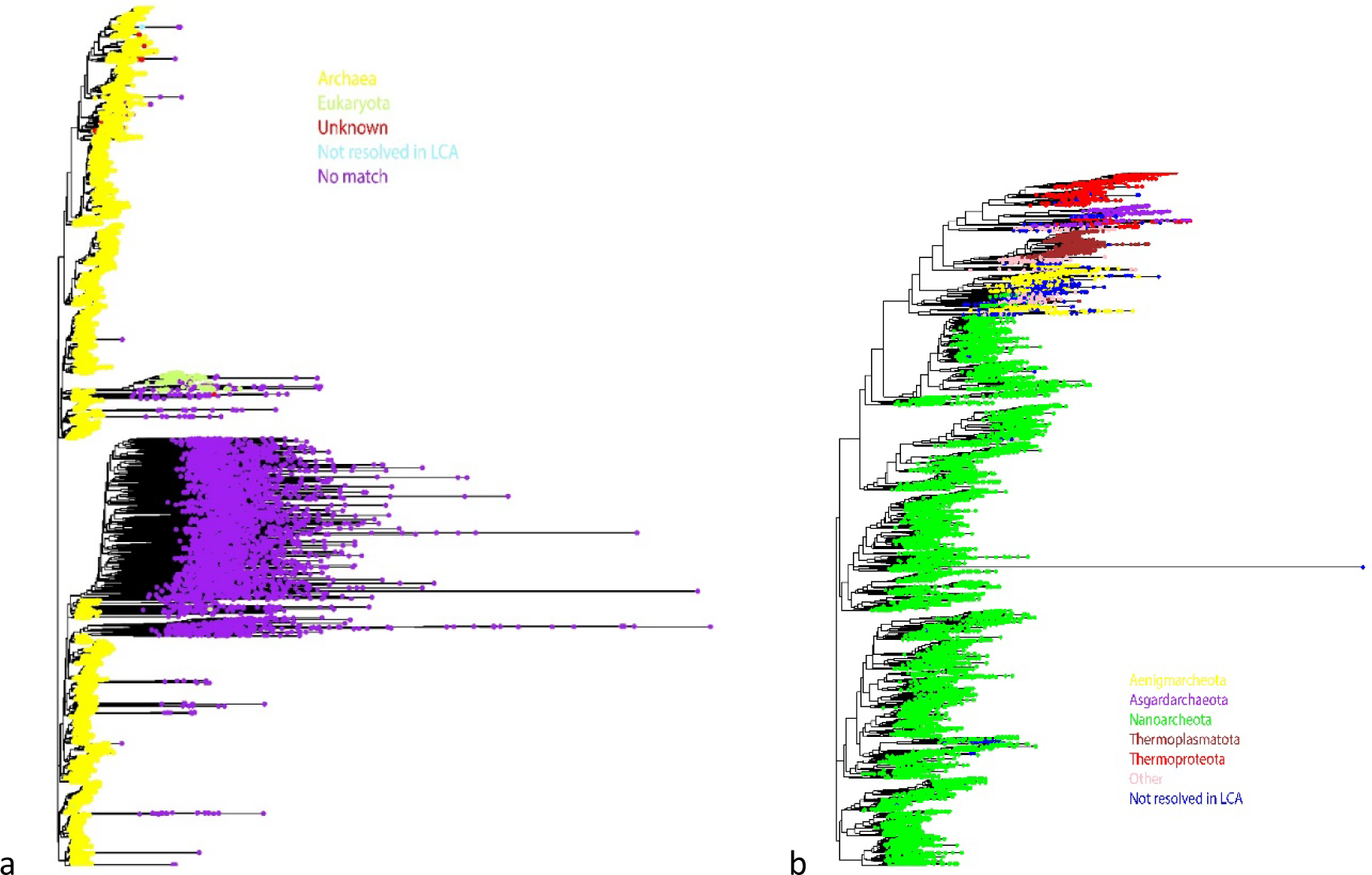
a) Maximum likelihood tree of all OTUs classified at domain level. “Unknown” indicates OTUs with matches to sequences identified as phylogenetically divergent Archaea by Karst et al. (2018) but not annotated by SINTAX. b) Maximum likelihood tree after removing OTUs identified as eukaryotes or Bacteria or without a database match. The 5 Dominant phyla are indicated. Other phyla are grouped together (pink) and Archaea where taxonomy is not resolved at phylum level by LCA are indicated in blue.

### Archaea taxonomic annotation performance of KSGP

Within LotuS2, we used the option “-keepUnclassified 0” to exclude those OTUs without a sequence similarity match in KSGP and the “-taxExcludeGrep” option to exclude eukaryote sequences and those with domain identified as “unknown”. This eliminated almost all of the long branches seen in Fig. 3a, leaving 11 451 OTUs, all but 9 of which had domain level assignments (dendrogram not shown, but very similar to Fig. 3b). 96.8% of these were assigned to phyla, 83.0% to class and 66.2% to order (Fig. 4a). Using SILVA and GTDB+ as references instead showed much lower assignment rates, except at phylum level: 94.6%/95% to phylum; 31.2%/33.9% to class and 15.8%/17.4% to order, respectively. The improvements in taxonomic annotation are 145%/166% and 280%/319% over GTDB+/SILVA at Class and Order level, respectively. KSGP also has many more high similarity matches than other databases, particularly for Altiarcheota, Iainarcheota, Nanoarcheota and OTUs not assigned to a phylum (Fig. 4b).

**Figure 4.**
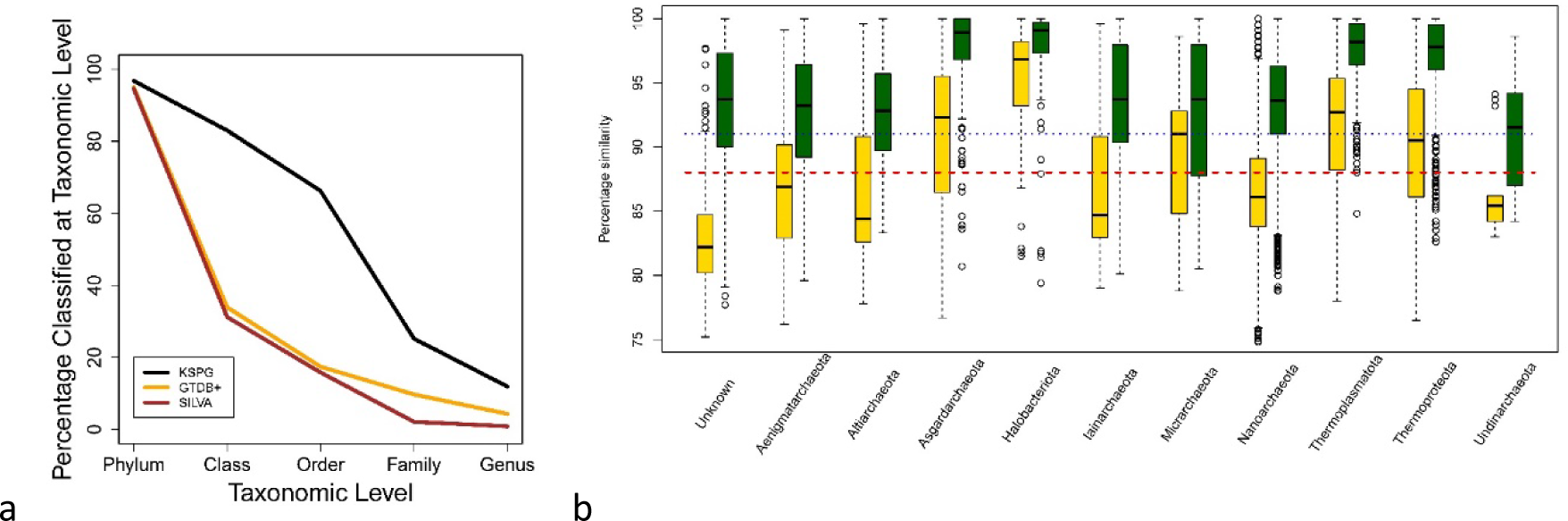
Taxonomic assignments. a) Percentages of archaeal OTUs with taxonomic assignment at levels from Phylum to Family based on three different databases – KSGP, GTDB+ and SILVA. *b)* Box plot of sequence similarities to closest match to individual OTUs in GTDB+ (yellow) and KSGP (green), broken down by Archaea phylum as assigned using KSGP. Blue dotted and red dashed lines indicate 91% and 88% sequence similarity, the cut-offs used within LotuS2 for order and class level assignments respectively. Phyla with fewer than 10 OTUs were assigned are excluded

For OTUs which are not assigned to phylum, the median sequence similarity of the nearest match increases from well below the 88% threshold for classification at Class level using GTDB+ to around the 93% threshold for assignment to families using KSGP. The level of annotation success for an individual OTU depends upon both the sequence similarity of the nearest matches and the level to which these matches are annotated. Although SILVA has a greater number of high-similarity matches than GTDB+, many of these have limited taxonomic information. By contrast, all sequences in GTDB are annotated to species level. As a result, LCA analysis based on GTDB+ has a slightly higher success rate than that based on SILVA (Fig. 4a).

We examine the taxonomic assignments of our three commonest OTUs to illustrate what lies behind these improvements. Within GTDB+, our most abundant OTU has 100% sequence similarity with *Nitrosopumilus* sp018263505 and *Nitrosopumilus cobalaminigenes* strain HCA1 (accession NR_159206) in NCBI nt. The second and third OTUs have 92% and 88% sequence similarity to their nearest matches in GTDB+, which are Thermoproteota MAGs (BIN-L-1 sp024650005 and PALSA-986 sp003164295 respectively) while their nearest matches in KARST (and thus KSGP) have 97.3 and 100% similarity respectively. KSGP classifies the first two OTUs to species level and the third to genus level, reflecting high confidence of the SINTAX taxonomic assignments of the KARST sequences incorporated into KSGP. This is entirely appropriate for OTU 1 but could be viewed as over-classification for the other two (see discussion).

KARST also includes information on the environment from which each sequence was retrieved. The great majority of OTUs have a closest match to a sequence from sediments and 45% of these are from one sample sd04, a muddy sediment collected from Limfjorden, Denmark (Table S2). There is an element of database coverage bias here, as the great majority of their archaeal sequences are from sediments and the largest number of these are from sample sd04. However, when standardised for database coverage our OTU sequences show significantly more matches than expected with many of Karst et al’s (2018) marine sediment samples, including sd04, and significantly fewer matches with freshwater sediments and an anaerobic digestor. Surprisingly, an estuarine sediment (sd10) also has significantly fewer matches in our data.

In our metabarcoding data, 79.6% of archaeal OTUs are assigned to the phylum Nanoarchaeota (Fig. 5) and of these, 69.5% to class Nanoarchaeota and 46.5% to order Woesearcheales. However, the relative abundance of these OTUs is, on average, lower than for those in other phyla, so although the most OTUs are from Nanoarchaeota, the phyla Thermoproteota and Thermoplasmatota are more prominent as a proportion of sequences (Fig. 5).

**Figure 5.**
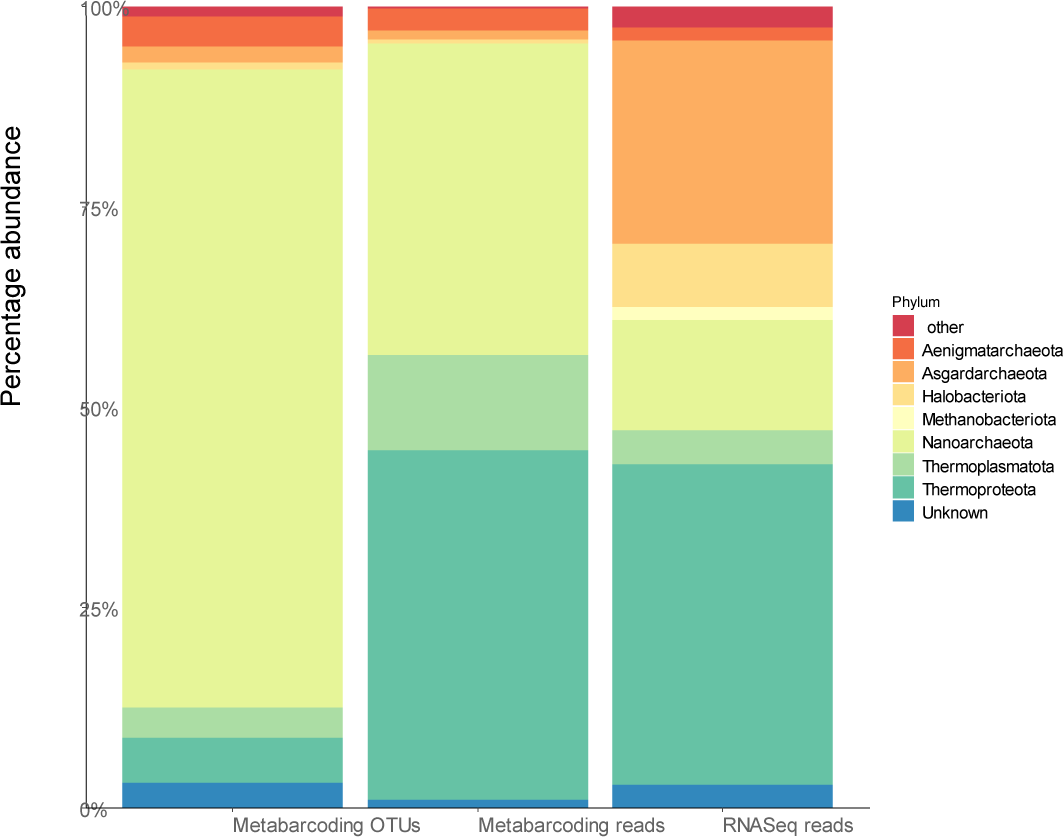
Phylum assignments of archaeal sequences from Estuarine sediment samples using KSGP, after excluding eukaryotes, bacteria and OTUs without a database match. Columns are metabarcoding data annotated by proportion of OTUs and proportion of reads, both of which are subject to PCR primer bias, and proportion of RNAseq reads.

### RNASeq results

Using a second dataset to distinguish the relative proportion of domains without PCR primer bias, we obtained 537,330 RNA reads, referred to as “RNAseq” data in the following. Ribosomal depletion protocols were not used, so most sequences represent rRNA, predominantly SSU and LSU sequences in approximately equal numbers (Turner et al., 2013). This avoids primer biases usually associated with amplicon sequencing experiments.

Archaea make up a rather small proportion of the community (448 reads) compared to bacterial (190,651) and eukaryotic (93,201) SSU sequences. In relative terms, approximately 0.23% of prokaryote RNA sequences and 0.16% of annotated sequences were of archaeal origin, lower than the 0.5% of all RNA sequences that we have previously reported for soil (Turner et al., 2013) or 0.8% of all PCR amplicons for boreal lakes (Juottonen et al., 2020) or 4% of prokaryote RNA sequences reported by Karst et al (2018, calculated from data in their Table S3). Thermoplasmatota make up a similar proportion of reads in the RNASeq data and metabarcoding sequences, but Nanoarchaeota are less abundant while Asgardarchaeota are more common (Fig. 5).

## Discussion

Curated reference sequence databases play a crucial role in metabarcoding and metatranscriptomics, having potentially major impacts on data quality of thousands of studies. Analysing amplicon sequences from sediments revealed significant shortcomings with current taxonomic databases to annotate and name Archaea. After removing apparent PCR artefacts, only 37% of our Archaea OTUs had a match in SILVA at > 88% sequence similarity expected within a Class. In addition, many of the sequences in SILVA and other databases are relatively poorly annotated because phylogenetic tree construction can be a significant challenge if only 16S rRNA gene sequence – or a part thereof – is available. There is only a small difference in the distribution of sequence similarities between our OTUs and either SILVA or GTDB+. However, all sequences in GTDB have a full taxonomic annotation is based on a large number of coding genes so both taxonomies and phylogenies will be more robust than those based on the 16S rRNA gene only. Further, GTDB is being regularly updated, whereas the most recent release of SILVA dates from 2019 (https://www.arb-silva.de/) amd at the time of writing the RDP website is no longer operational (http://rdp.cme.msu.edu/). Greengenes was last updated in 2019, although has been superseded by https://greengenes2.ucsd.edu/) in 2023. For all these reasons, GTDB offers a more future proof basis for taxonomies and phylogenies.

GTDB SSU sequences are naturally much more limited in number, as GTDB incorporates only near complete genomes/MAGs. They do have taxonomic assignments down to the level of species that are based on a much larger evidence base than SSU sequences alone. However, even the sparse archaeal GTDB rRNA gene sequences required further polishing from technical artifacts and obvious misclassifications. Without this, large fractions of OTUs without close homologs in the database could not be resolved at domain level (details not shown). Contaminant sequences could have equally strong impacts on classifications at lower taxonomic levels. There is growing recognition of the challenge posed by contamination of genome sequences (Arkhipova, 2020; Orakov et al., 2021) and the processing of genomes by GTDB includes checks for contaminants (Parks et al., 2021), but some additional screening of the SSU sequences is still required. The SATIVA subset of GTDB (Quast et al., 2012) has a similar objective. But as well as removing misclassified sequences it also excludes sequences that are short or contain ambiguities, which is the case for the majority of archaeal MAG derived genomes in GTDB. Probably due to a combination of strict filtering and reliance on older database versions, SATIVA had a substantially poorer coverage of our OTUs and hence led to poorer taxonomic assignments and the same was also true of Greengenes2. As a result, neither can be recommended as sole taxonomic reference for metabarcoding studies of Archaea. GTDB should be supplemented with a eukaryote SSU database, because archaeal and eukaryotic SSU sequences can be more similar to each other than to bacterial SSU sequences (c.f. Eme et al., 2023; Zaremba-Niedzwiedzka et al., 2017) and hence archaeal primers sometimes amplify eukaryotic SSUs. We included eukaryote sequences in KSGP to avoid mismapping eukaryote sequences to archaeal rRNA sequences. For the environments that we have sampled, PR2 has a good coverage of eukaryote sequences that are amplified, although for non-marine environments other databases, such as the eukaryote sequences in SILVA, might be preferred.

A further challenge is that there are many archaeal lineages at Class, and even Phylum, level that are not included within either SILVA or GTDB (Karst et al., 2018 and see Figs. 1 and 2), although common OTUs are more likely to be successfully annotated than rare ones. Recent revisions to Archaea phylogeny even at phylum level indicate that their phylogenetic diversity remains to be fully characterised, certainly at the level of class and perhaps at phylum level (Liu et al., 2021; Rinke et al., 2021). This is supported by Karst et al.’s identification of eight deeply branching lineages of Archaea.

We show that this lack of very close sequence matches in databases can be partially overcome by using the consistent GTDB taxonomy to annotate large sets of environmental rRNA gene sets such as that published previously by Karst et al. or Archaea sequences from SILVA. Annotating using a k-mer matching algorithm such as SINTAX (Edgar, 2016a) rather than the phylogenetic tree based approaches of SATIVA (Lundin & Andersson, 2021) or Greengenes2 (Daniel McDonald et al., 2023) allows us to incorporate information from incomplete SSU sequences. This leads to a substantially higher level of classification success than obtained using either SATIVA or Greengenes2. KSGP, the modified GTDB+, supplemented with these additional sequences, allows us to identify the great majority of archaeal OTUs to phylum level and beyond, while also providing a strategy to exclude PCR artefacts and bacterial OTUs. It also allowed us to substantially increase the number of annotated OTUs to class and order level. After PCR artefacts are excluded, KARST contains sequences with > 90% sequence similarity to more than 80% of our OTUs suggesting that the great majority of our OTUs are in the same class, order or even family as particular sequences in KARST, based on sequence similarity within higher level taxa (Yarza et al., 2014). Our approach does not, for example, successfully annotate all of Karst et al.’s deeply branching Archaea even at domain level, but this will improve as genomes from deeply branching lineages are added to GTDB over time. The use of an LCA strategy within LotuS2 (Ozkurt et al., 2022; Yarza et al., 2014) will likely remove false positive assignments to KSGP, but it is likely that taxonomic annotations within poorly characterised parts of the phylogeny will be less robust than those achieved directly with GTDB+. In consequence, we can be less confident in the names that we assign to these taxa. The assignments should be interpreted as indicating the closest match amongst taxa that *are* robustly placed on a phylogenetic tree rather than as a definitive taxonomic assignment. It may be wise to preface them with “c.f.” (sensu Lucas, 1986) to indicate this and, for common OTUs, examine in more detail levels of sequence identity with the closest matches in both GTDB+ and KSPG, as illustrated above for our three commonest OTUs. Such classifications are, nevertheless, very useful to guide more detailed study of the components of the community and it will be easier to clarify these phylogenetic relationships using the near full length sequences from KARST rather than shorter amplicon sequences. In our case, the relatively high diversity found in Nanoarchaeota, and order Woesearcheales in particular is intriguing, and the much higher sequence similarities for this phylum in KSGP than in GTDB+ (Fig. 4) suggests that phylogenetic diversity at the level of class and order within this phylum is not yet captured by GTDB and the MAG-centric studies represented in this databse. The same strategy that we have used with the Karst data could be used to add other near full-length sequences to a custom database or to provide an independent check on taxonomic assignments in existing databases. We have chosen to use a threshold probability of 0.8 when annotating the SILVA and Karst et al sequences, but a more stringent probability may be preferred by some.

The SSU1ArF/SSU520R primer pair was designed to amplify as broad a range of Archaea, but there is likely yet phylogenetic diversity that is still being uncovered (Tahon et al., 2021; Yang et al., 2022; Zhang et al., 2020) and the RNASeq data suggest this primer pair may generate PCR biases against Asgardarcheota and in favour of Nanoarcheota, (or that the former are metabolically less active while the latter are more active than other Archaea). Therefore, we recommend that RNAseq is carried out in conjunction with metabarcoding to give a less biased taxonomic representation of a community and the relative importance of Archaea, Bacteria and eukaryotes (c.f. Turner et al., 2013). Full length SSU RNA sequences would be the gold standard for this RNASeq work, as used in Karst et al. (2018), but their methods are technically challenging and relatively time consuming (Eisenstein, 2018). For quantifying absolute abundance, we have shown that library preparation from total RNA using a commercial RNA kit allows us to assess the relative abundance of Bacteria and Archaea, yielding more SSU sequences than metagenomic and metabarcoding approaches (c.f. Eisenstein, 2018; Eloe-Fadrosh et al., 2016).

## Conclusions

The coding-gene based taxonomy established by GTDB provides a robust foundation for annotation of Archaea metabarcoding data. GTDB alone gives a small improvement in the taxonomic annotation of our archaeal data, but a much greater improvement is obtained when GTDB is used to annotate SILVA archaea sequences and a collection of near full length rRNA sequences to create the KSGP database. KSGP is also likely to facilitate improved annotation of bacterial metabarcoding data (see supplementary material). Most of this improvement comes from including the Karst et al. (2018) sequences, indicating that future attempts to expand database coverage should focus on direct rRNA sequencing rather than on PCR amplicons with their associated primer biases. This is likely to be particularly useful for environments that are not well covered in the Karst et al. (2018) data.

### Box 1

- A recommended strategy

Based on the analyses presented above, we recommend the following approach to the analysis and taxonomic annotation of archaeal metabarcoding data:

1. Use UPARSE (or another similar approach) to generate 0.97 similarity radius OTUs, unless there are *a priori* reasons for focussing on sub-specific patterns and using ASVs/zOTUs.
2. Examine the extent to which GTDB, KSGP (and SILVA if desired) cover the phylogenetic diversity represented by these OTUs using the approaches demonstrated in Figs. 1 and 2.
3. Use GTDB, combined with a curated database of eukaryote sequences such as PR2, to carry out robust taxonomic assignments based on cultivated strains and MAGs and identify OTUs assigned to Eukarya or Bacteria, or without a database match.
4. Use KSGP to provide taxonomic assignments based on environmental SSU sequences annotated with the GTDB taxonomy, recognising that assignments that are substantially better than those using GTDB alone should be prefaced by c.f. and that phyla, classes and orders that are not resolved in this process are likely to represent undescribed taxa, which can be investigated further using additional option (a) below.
5. Examine OTUs without database matches, carry out Blast searches against NCBI nt on the commonest and compare ML trees with and without these OTUs. If substantial numbers of common (low numbered) OTUs are being removed, some of the additional options below should be considered to understand these in more detail.
6. Carry out downstream analyses using the OTU table and phylogenetic tree with non-matched OTUs, Bacteria and Eukarya omitted.

Additional options.

a. For common OTUs without a high similarity match in GTDB, use the OTU sequence to search for matches in NCBI nt or other databases using the less stringent BlastN strategy. Eukaryote OTUs, for example, will be identified by this. If there are close matches in KSGP, searches can be carried out using these longer sequences to improve taxonomic resolution and they could also be used in constructing a phylogenetic tree.
b. For common OTUs with a relatively poor match in GTDB, examine taxonomic assignments and the environmental source of closest matching sequences in NCBI nt and Karst
c. Total RNAseq using short reads provides estimates of the relative abundance of taxa without PCR bias so is a very valuable addition to metabarcoding. Long rRNA sequences obtained using a linked read approach similar to that used by Karst et al or PacBio reads of cDNA prepared from total RNA would enable the approach that we have used here for environments where equivalent data to Karst et al are not available.

## Acknowledgements

Abdullah Aleidan was supported by King Saud University and Solomon Udochi by the Commonwealth Scholarship Commission. MB was supported by the Swedish Research Council (Vetenskapsrådet; Grant 2021–03724). JF was supported by the UKRI Biotechnology and Biological Sciences Research Council Norwich Research Park Biosciences Doctoral Training Partnership, BB/T008717/1. The bioinformatic analysis was carried out on the High Performance Computing Cluster supported by the Research and Specialist Computing Support service at the University of East Anglia. F.H. was supported by European Research Council H2020 StG (erc-stg-948219, EPYC) and by the Biotechnology and Biological Sciences Research Council (BBSRC) Institute Strategic Programme (ISP) Food Microbiome and Health BB/X011054/1 and its constituent project BBS/E/F/000PR13631; Earlham ISP BBX011089/1 and its constituent work package BBS/E/ER/230002A.

## Data Accessibility

The KSGP database and the cleaned GTDB database are available at ksgp.earlham.ac.uk. Unprocessed DNA sequences and sequences of OTUs are available as ENA accession PRJEB65254.

## Author Contributions

AG conceived the study, performed the data analyses and wrote the paper; AA; CS and SCU collected the data; FH developed the LotuS2 pipeline; adapted it to the needs of this study; and wrote the paper; JF contributed software development and MB contributed to writing the manuscript.

## Supplementary Material

Table S1. Sequences removed from PR2 and GTDB as assigned to incorrect domain

Table S2. Numbers of OTUs with closest match to RNA sequence obtained from individual environmental samples in the Karst et al. (2018) data set

Appendix 1. Comparison of KSGP with GreenGenes2, including:

Figure S1. Rank order plots of highest similarity to each OTU in the GreenGenes2 full database and backbone; KSG and GTDB+ for bacterial and Archaea OTUs.

**Table S1.**
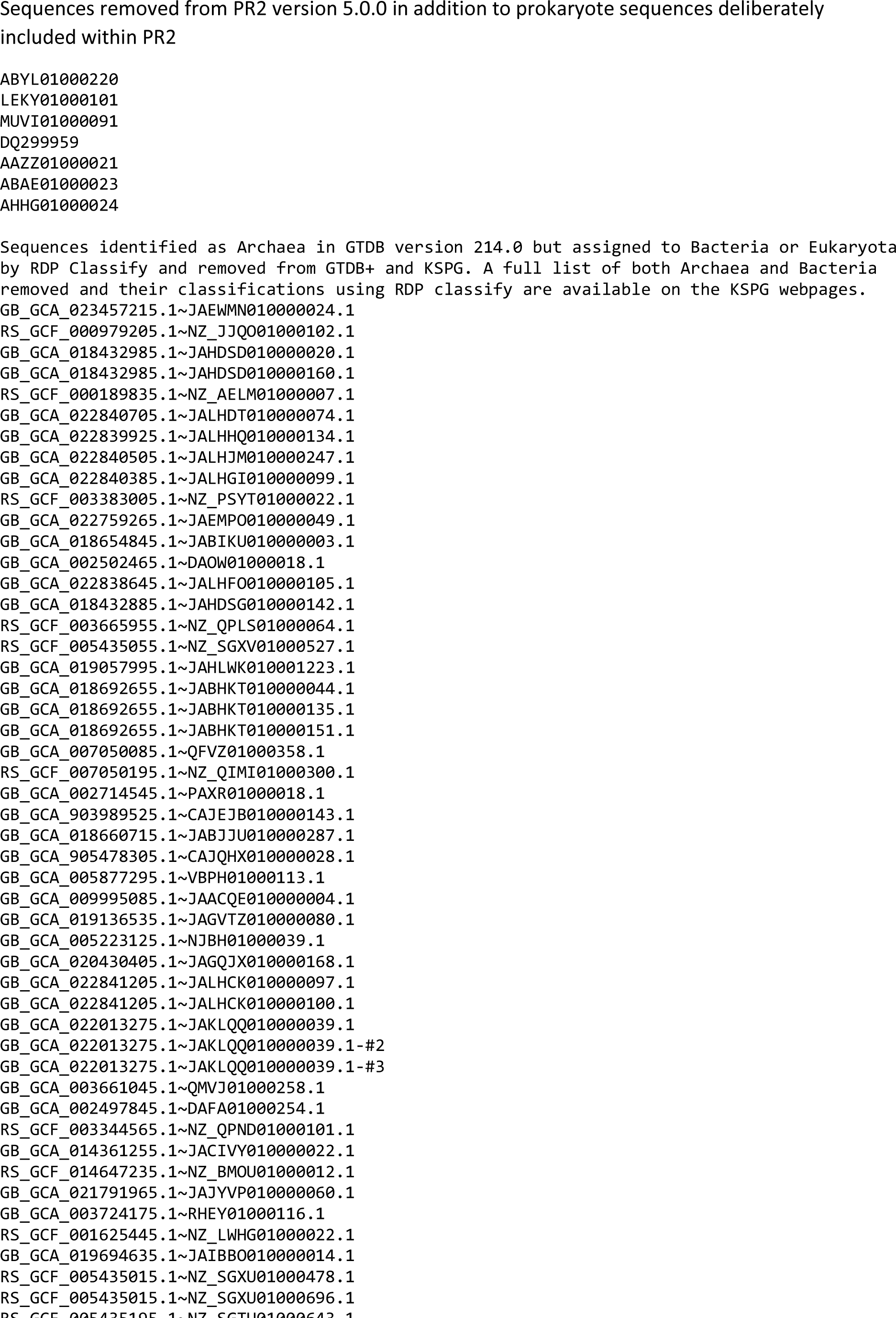
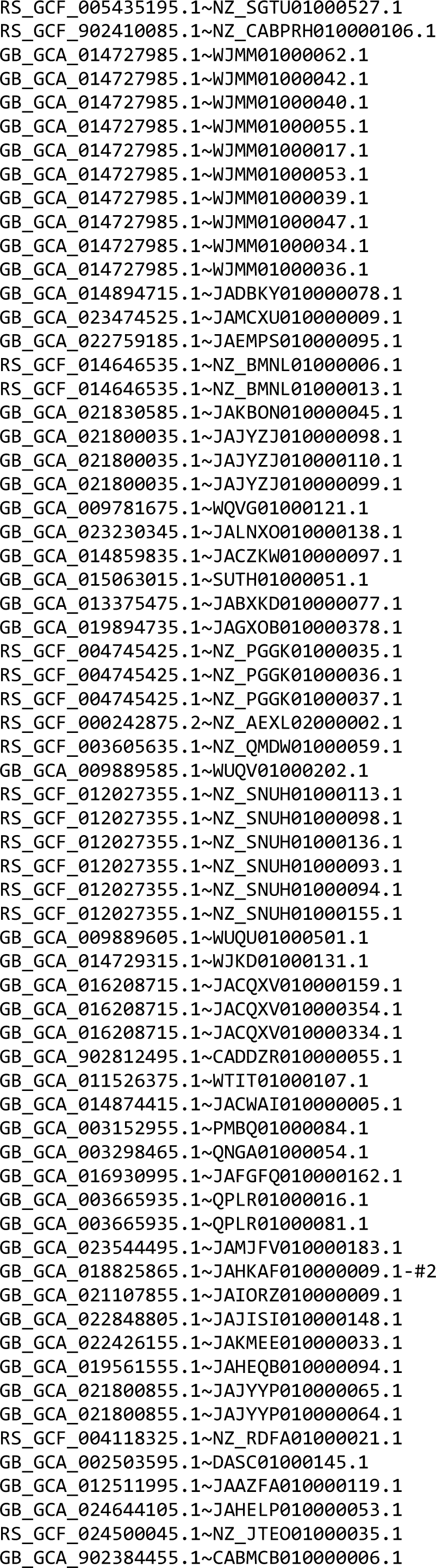
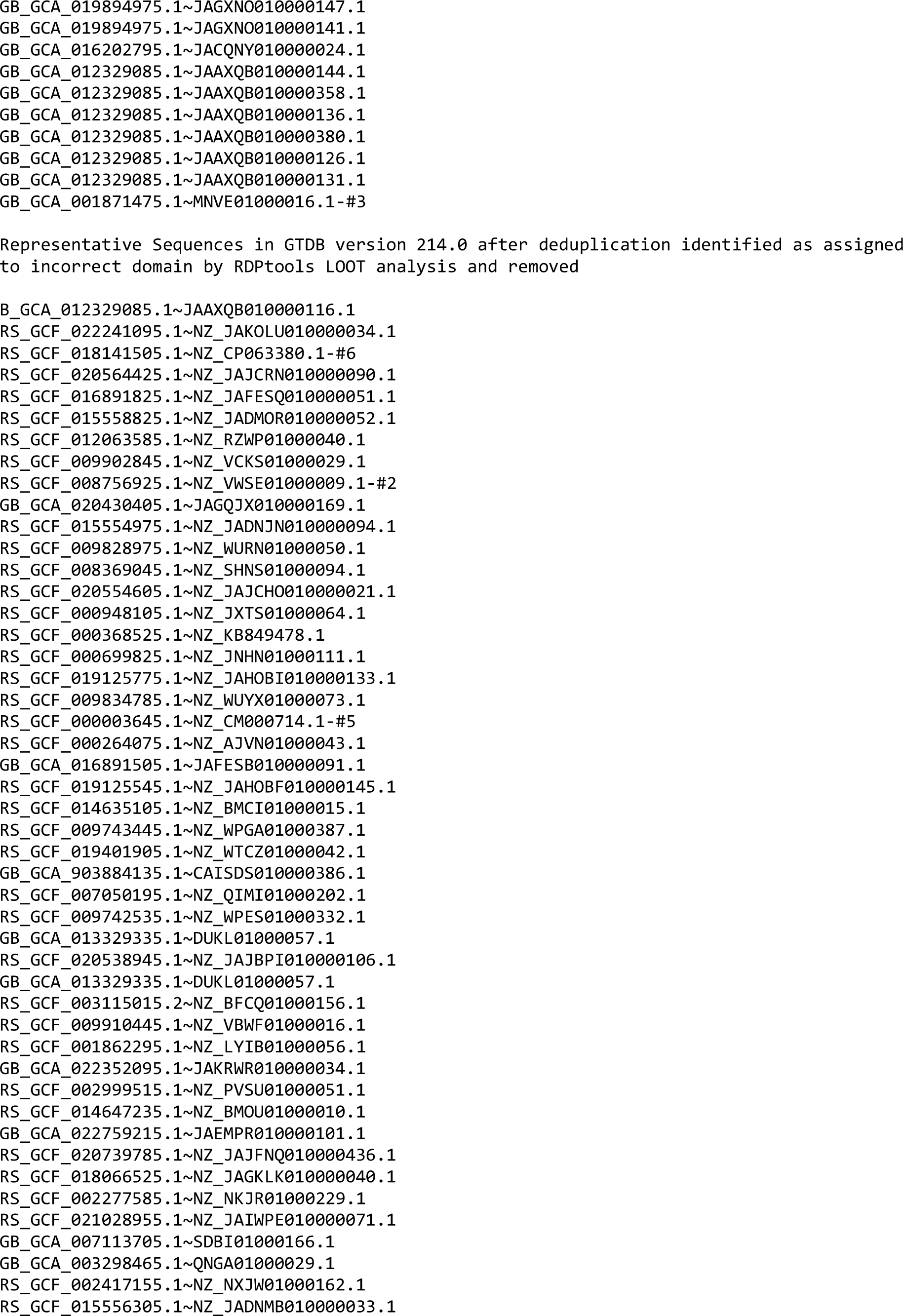
Sequences removed from PR2 and GTDB as assigned to incorrect domain

**Table S2.** Numbers of OTUs with closest match to RNA sequence obtained from individual environmental samples in the Karst et al. (2018) data set

| **Karst sample** | **Archaea 16S**<br>**sequences from that**<br>**Sample in Karst** | **Number of OTUs**<br>**with closest match**<br>**from Karst sample** |
| --- | --- | --- |
| ad01A – Anaerobic digester | 1654 | 12 |
| as01A – Activated Sludge | 46 | 5 |
| fw01A – Freshwater | 7 | 2 |
| gw01A – Ground Water | 131 | 51 |
| hg01B – Human Gut | 1652 | 5 |
| *Marine and Estuarine*<br>*Sediments* |  |  |
| sd01A | 29 | 9 |
| sd01B | 2161 | 1353 |
| sd02A | 12 | 6 |
| sd02B | 1065 | 625 |
| sd03A | 55 | 16 |
| sd03B | 601 | 327 |
| sd04A | 468 | 177 |
| sd04B | 12646 | 5086 |
| sd09A | 1668 | 508 |
| sd10A | 5327 | 469 |
| sd10B | 10393 | 1599 |
| sd12A | 928 | 491 |
| *Freshwater sediments* |  |  |
| sd05A | 10235 | 323 |
| sd06A | 4093 | 250 |
| sd07A | 802 | 78 |
| sd08A | 1512 | 121 |
| sd11A | 5115 | 116 |
| *Soil* |  |  |
| so01A | 5 | 1 |
| so01B | 14 | 33 |
| so02A | 601 | 40 |

## Appendix 1

### More detailed comparison of KSGP with Greengenes2 for both Archaea and Bacteria

The Greengenes2 database (McDonald et al., 2023) is based on an apparently similar approach to KSGP, so its relatively poor coverage of our OTUs merits further discussion. For Archaea, the Greengenes2 backbone produces lower similarity matches than GTDB+ (Fig. S1), presumably because incomplete SSU sequences have been removed, impacting coverage of Archaea more markedly than for bacteria. The same phenomenon is seen with the SATIVA subset of GTDB. There may also be some contribution due to incomplete coverage of Eukaryota in Greengenes2. The inclusion of amplicons in the full Greengenes2 database has minimal effect on coverage of Archaea. The latter is likely to be a consequence the great bulk of these amplicons being short bacterial sequences – the average length of sequences in the full Greengenes2 database is 145.4 bp. KSGP performs substantially better than either the Greengenes2 backbone or the full Greengenes2 database.

For bacteria, the Greengenes2 backbone has more comprehensive coverage than GTDB+. But again, the addition of a very large number of amplicons to the full Greengenes2 database makes only a small improvement to database coverage and KSGP performs substantially better than both.

**Figure S1.**
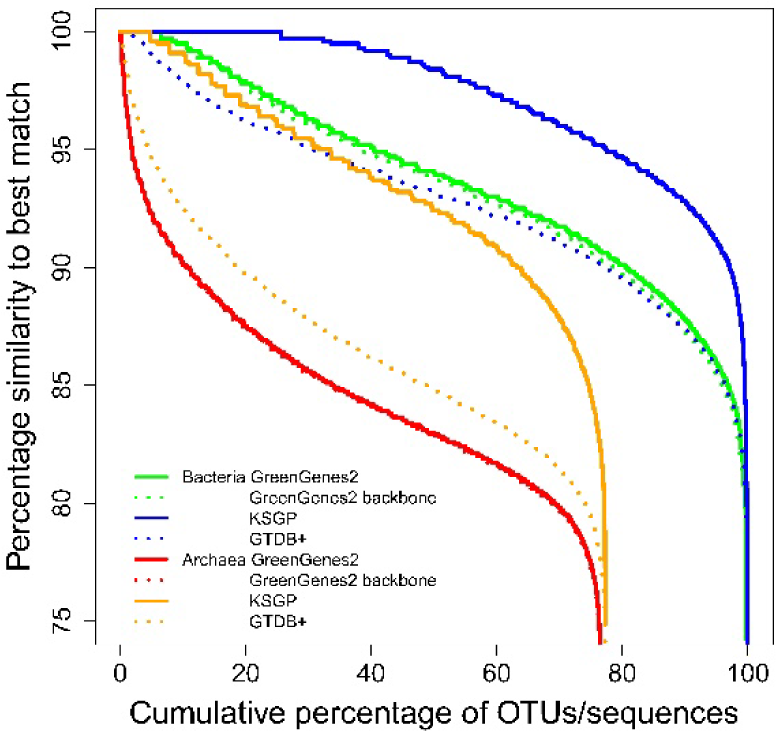
Rank order plots of highest similarity to each OTU in the GreenGenes2 full database and backbone; KSGP and GTDB+ for bacterial and Archaea OTUs. Similarities were generated using USEARCH_local (Edgar, 2010) rather than UBLAST due to the high memory requirement of carrying out low stringency searches on the full Greengenes2 database. Lines for Greengenes2 backbone are almost identical to those for the full database

